# Joint Biophysical Modeling of Paired Single-Cell RNA and Protein Measurements

**DOI:** 10.1101/2025.11.14.688548

**Authors:** Catherine Felce, Meichen Fang, Lior Pachter

## Abstract

Surface protein measurements can supplement gene expression information from single-cell RNA sequencing to provide a more complete assessment of cell identity and function. Recently developed multiomic assays facilitate such measurements, and can, in principle, be utilized to understand the dynamics of transcription and translation. We develop a framework for biophysical modeling of transcription jointly with translation from single-cell data, along with a suitable technical noise model for sequencing data. We demonstrate its efficacy using simulations, and illustrate how it can be useful in practice with 10x multiomic data. Our proof-of-principle highlights the potential for jointly modeling transcription and translation as data quality and measurement accuracy improves.

## 1 Background

Although messenger RNA has historically been used as a proxy for protein expression (Junker et al. 2014; Fu et al. 2007), many studies have highlighted the role of post-transcriptional regulation in decoupling RNA and protein levels (Wei et al. 2015; Franks et al. 2017), especially in embryogenesis and other developmental processes (Kuersten and Goodwin 2003). The poor correlation between mRNA and protein counts across the genome has been extensively investigated (Maier et al. 2009), with biologists calling for principled modeling (McManus et al. 2015; Greenbaum et al. 2003), including technical noise models (Liu et al. 2016), to shed light on these discrepancies. While differentially expressed RNA has been shown to correlate more strongly with its associated proteins than non-differentially expressed RNA (Koussounadis et al. 2015), the range of correlations for individual genes is still broad (*r* = −0.95 to 0.94) and differentially expressed proteins have been shown not to reliably correlate with differentially expressed RNA (Huber et al. 2004; Ideker 2001).

Given that combined RNA expression and proteomic data are helpful for analyzing the immune response (Wu et al. 2020) and bacterial growth (Nobori et al. 2020), there is a clear benefit to investigating the separate regulatory contributions of transcription and translation using multiomic data. Recognizing this, statistical tools have been created to disentangle RNA and protein-level regulation using time-course data (Teo et al. 2014), and studies have confirmed that RNA and protein-level regulation are both important, with some genes being regulated in opposite directions simultaneously (Cheng et al. 2016). Cross-species studies have also highlighted the potential decoupling of mRNA and protein levels, and have tried to unravel the effects of transcriptional, translational, and post-translational mechanisms on expression divergence (Wang et al. 2018).

However, time-course or cross-species measurements are difficult to obtain, and increasingly ubiquitous single-cell data offer the opportunity, in principle, to study the dynamics of transcription and translation in a complementary way. We propose a biophysical model based on single-cell multiomic data that can simultaneously describe transcriptional and translational mechanisms and untangle their contributions to controlling protein expression.

Single-cell resolution RNA measurements have already enabled biophysical modeling of nascent and mature RNA and the inference of transcriptional rate parameters (Gorin et al. 2023). In that setting, the modeling relies on multimodal datasets, specifically spliced and unspliced counts measured from mature and nascent RNAs (Gorin et al. 2022). The inclusion of chromatin information from ATAC-seq has also been considered (Felce et al. 2024). In this work we extend these approaches to include protein count data.

High quality single-cell protein quantification, including isoform differentiation, has become available with single-cell western blotting (Hughes et al. 2014). Since 2017, with the advent of high-throughput simultaneous proteomic and transcriptomic measurements in single-cells with REAP-seq (Peterson et al. 2017) and CITE-seq (Stoeckius et al. 2017), principled biophysical modeling of the joint modalities is now within reach.

Although models combining RNA dynamics with constitutive protein translation of mature RNA transcripts have been suggested (Singh and Bokes 2012), we solve numerically for the probability distribution for such a system and demonstrate that this can be used to reliably infer biophysically meaningful parameters on real datasets. Our approach, summarized in Figure 1, provides a proof of principle for fitting biophysical models to multiomic data.

**Fig. 1:**
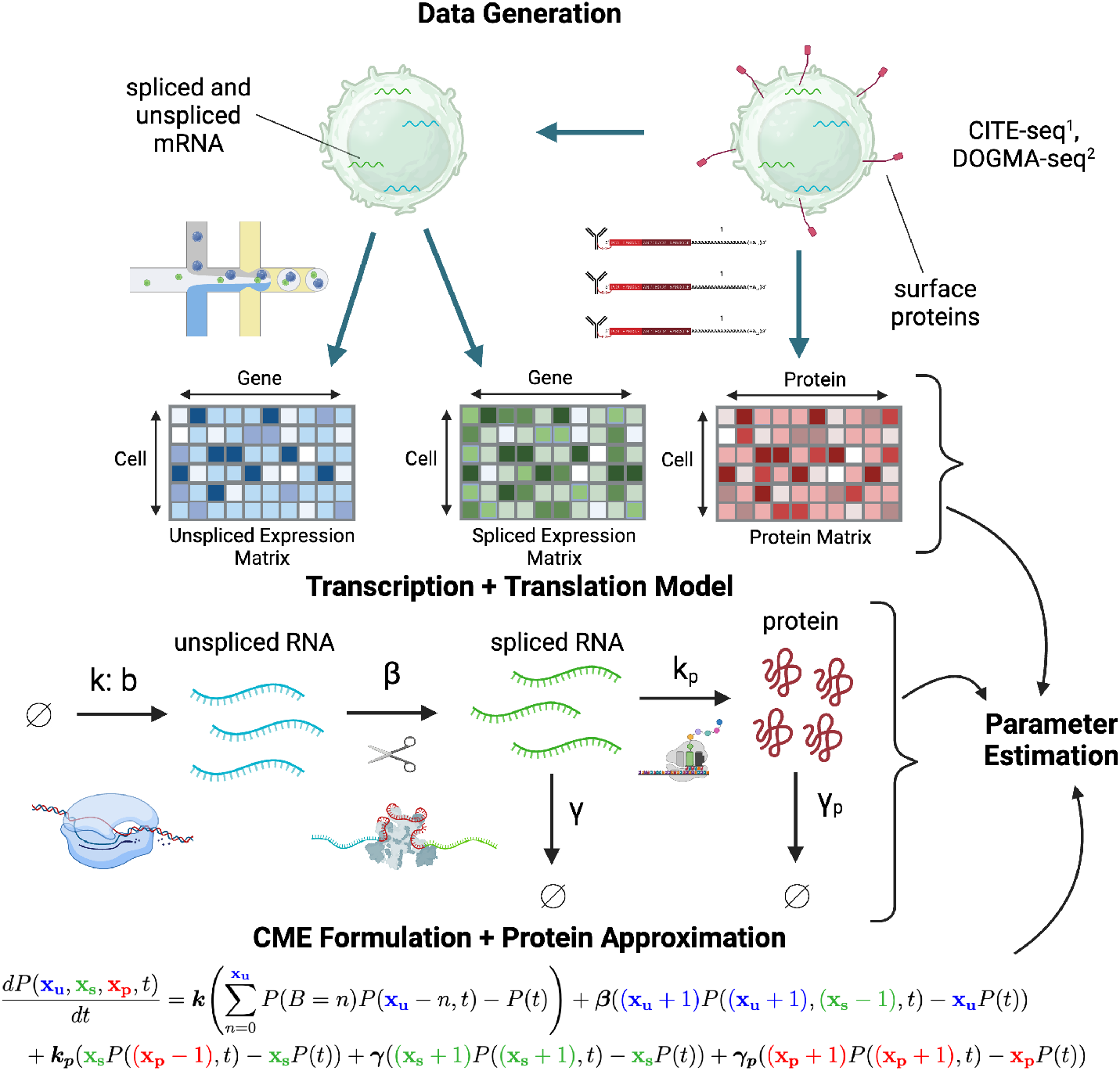
Fitting a joint biophysical model for RNA and protein. Single-cell transcriptomics and surface protein expression can be simultaneously measured using methods such as ^1^CITE-seq (Stoeckius et al. 2017) and ^2^DOGMA-seq (Mimitou et al. 2021). Transcriptomic data can be used to generate unspliced and spliced count matrices. We assume a bursty transcription and constitutive translations with constitutive splicing and degradation. We derive the chemical master equation (CME) following Singh et al. (Singh and Bokes 2012) and apply continuous approximation to protein when necessary. The resultant joint probability distribution of unspliced mRNA, spliced mRNA and protein is solved numerically and used for parameter estimation.

## 2 Results

### 2.1 Model

We consider the bursting limit of the telegraph model as in Singh and Bokes (Singh and Bokes 2012), with RNA processing from nascent to mature transcripts at a rate *β*, and translation rate per RNA molecule *k*_*p*_ (see Table 1 for all parameter identifications). This system can be summarized in the following reactions.

**Table 1:** Notation for the joint biophysical model, and expressions used for numerical solving of the steady-state probability distribution over unspliced, spliced and protein counts.

| Symbol | Meaning |
| --- | --- |
| $k \in \mathbb{R}_+$ | Burst frequency |
| $B \in \mathbb{N}$ | Burst size |
| $b \in \mathbb{R}$ | Mean burst size |
| $\beta \in \mathbb{R}$ | Splicing rate of unspliced transcripts |
| $\gamma \in \mathbb{R}$ | Decay rate of spliced transcripts |
| $k_p \in \mathbb{R}$ | Translation rate of spliced transcripts |
| $\gamma_p \in \mathbb{R}$ | Decay rate of proteins |
| $X_u \in \mathbb{N}$ | Unspliced RNA copy number |
| $X_s \in \mathbb{N}$ | Spliced RNA copy number |
| $X_p \in \mathbb{N}$ | Protein copy number |
| $P(x_u, x_s, x_p, t)$ | Probability density of state $(x_u, x_s, x_p) \in \mathbb{N}^3$ at time $t$ |
| $G(z_u, z_s, z_p, t)$ | Generating function (GF) of $P(x_u, x_s, x_p, t)$ |
| $\phi(u_u, u_s, u_p, t)$ | Factorial-cumulant GF $\log G(u_u + 1, u_s + 1, u_p + 1, t)$ |
| $F(z_u) = \frac{1}{b+1-bz_u}$ | Generating function of $B$, i.e., $\sum_{n=0}^{\infty} z_u^n P(B = n)$ |
| $M(u_u) = \frac{bu_u}{1-bu_u}$ | Transformation of the burst PGF, i.e., $M(u_u) = F(1 + u_u) - 1$ |

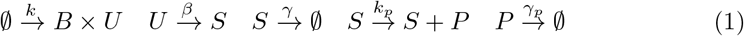

As in Singh et al. (Singh and Bokes 2012), we define the probability mass function

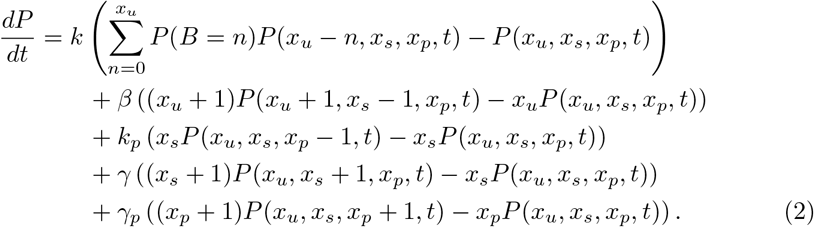

We then use generating function methods to calculate the stationary distribution. As in (Gorin et al. 2022; Gans 1960), we define the probability generating function, *G*, via

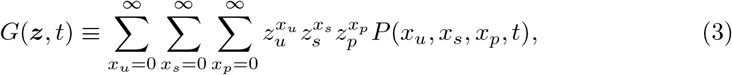

where ***z*** represents the vector of arguments (*z*_*u*_, *z*_*s*_, *z*_*p*_). We then define *ϕ* via

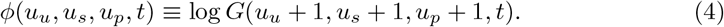

Using the method of characteristics (see the Supplementary Information, S1), we arrive at the following system of equations:

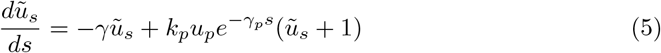

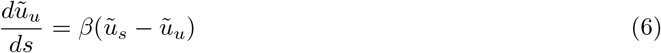

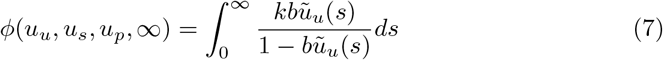

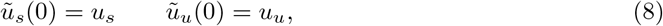

which can be solved numerically (see the Supplementary Information, S2).

On top of the biological model, which is summarized in Figure 1 **Transcription + Translation Model**, we also incorporate a technical noise model to account for noise in the sequencing process following (Gorin et al. 2025), with both RNA and protein counts assumed to undergo Poissonian sampling, whose parameters are optimized via grid search.

### 2.2 Simulation

First, we use simulated data to assess the accuracy of estimated parameters under our model. We simulated 100 genes by first drawing their true parameters from a biologically plausible range, then generating steady-state data using the Gillespie algorithm. The application of our mean-expression filter resulted in a final set of 46 genes. We applied our inference framework to these 46 simulated genes and were able to recover all of the parameters in the full, joint model with unspliced, spliced and protein counts. The inference accuracy is shown in Figure 2. Parameter estimates were generally accurate, except for two outliers whose protein degradation rates converged to the optimization lower bound. The two outliers reside in a regime of low mRNA and high protein expression, where absolute parameter values are less identifiable, and only the ratio of translation to protein degradation was correctly inferred. The simulation results suggest that we should exclude parameter estimates found at the optimization bounds when fitting to real datasets.

**Fig. 2:**
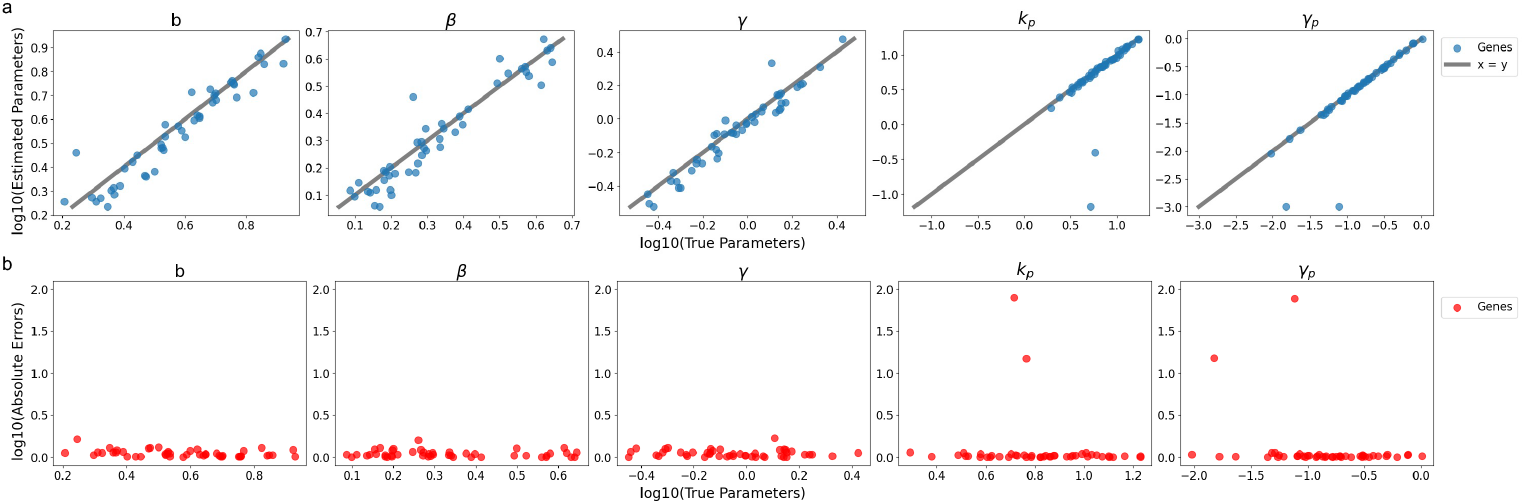
Inference accuracy on simulations. a) Estimated parameters versus true parameters b) Absolute errors versus true parameters.

### 2.3 Fits to Real Data

Given the simulation results, we first sought to apply the full model to the unspliced RNA, spliced RNA and protein counts from real datasets. However, we observed that unspliced counts are generally low and noisy, making it difficult to identify genes with high expression across all modalities. Therefore, we asked whether protein and spliced RNA counts are informative enough for inference. We refit the model using only the spliced mRNA and protein counts from the 46 simulated genes and found all parameters to be identifiable, though with a loss of accuracy in the splicing rate estimates (Figure 3). As we are more interested in the kinetic parameters related to translation, we decided to fit our model on the protein and spliced RNA counts of real datasets for now. The full model can be used when data quality permits in the future.

**Fig. 3:**
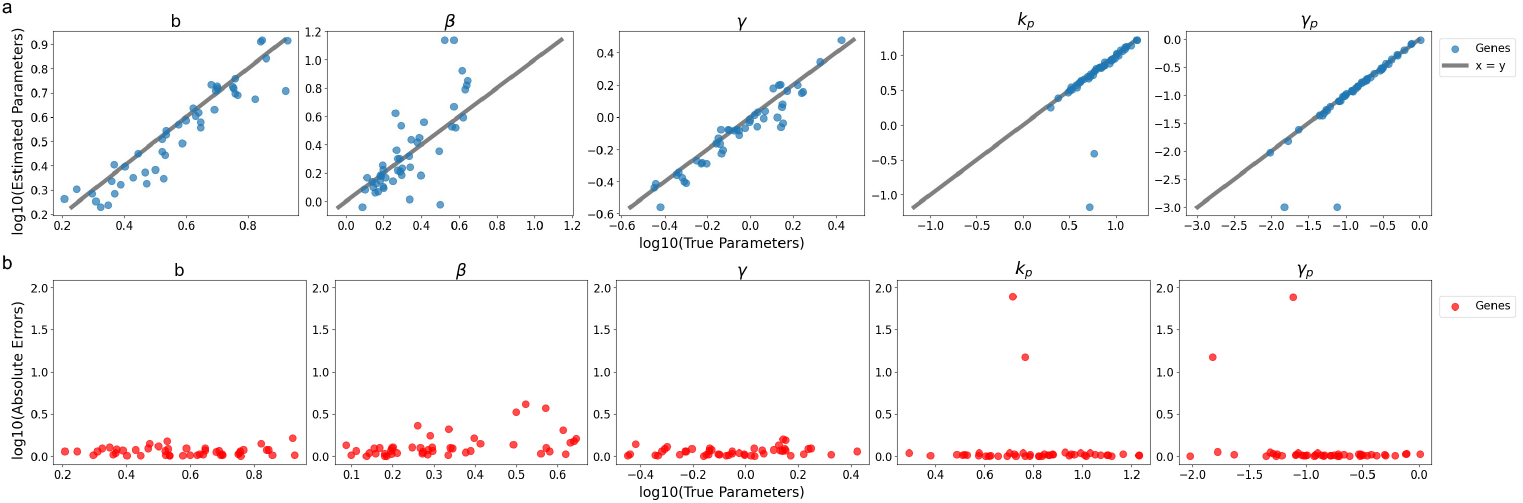
Inference accuracy with only spliced mRNA and protein counts on simulations. a) Estimated parameters versus true parameters b) Absolute errors versus true parameters.

In particular, we fit our model on the protein and spliced RNA counts from the following human PBMC datasets:

- 10k Human PBMCs Stained with TotalSeq™-B Human TBNK Cocktail, Chromium GEM-X Single Cell 3’ Universal 3’ Gene Expression dataset analyzed using Cell Ranger 8.0.0 (2024, March 13) (10x Genomics 2024)
- 10K Human PBMCs, Gene Expression with a Panel of TotalSeq™-B Antibodies, analyzed using Cell Ranger 3.0.0, (2018, November 19) (10x Genomics 2018)

We processed the raw transcript reads using kb-python (kallisto and bustools) to obtain spliced and unspliced RNA counts. We used the cell-by-protein count matrix provided by 10x Genomics. For protein complexes with subunits, we assumed that the presence of the complex indicates the presence of all constituent subunits, so we chose from among the subunits when mapping to RNA transcripts. More details on data processing can be found in the Supplementary Information (S3.1).

In Figure 4, we show the fits of our model to a few marker genes. The technical sampling parameters to which these fits correspond, and the optimal fitted biophysical parameters, are given in the Supplementary Information (S3.2). The cells were subsetted to monocytes and T cells for the 2024 and 2018 datasets respectively.

**Fig. 4:**
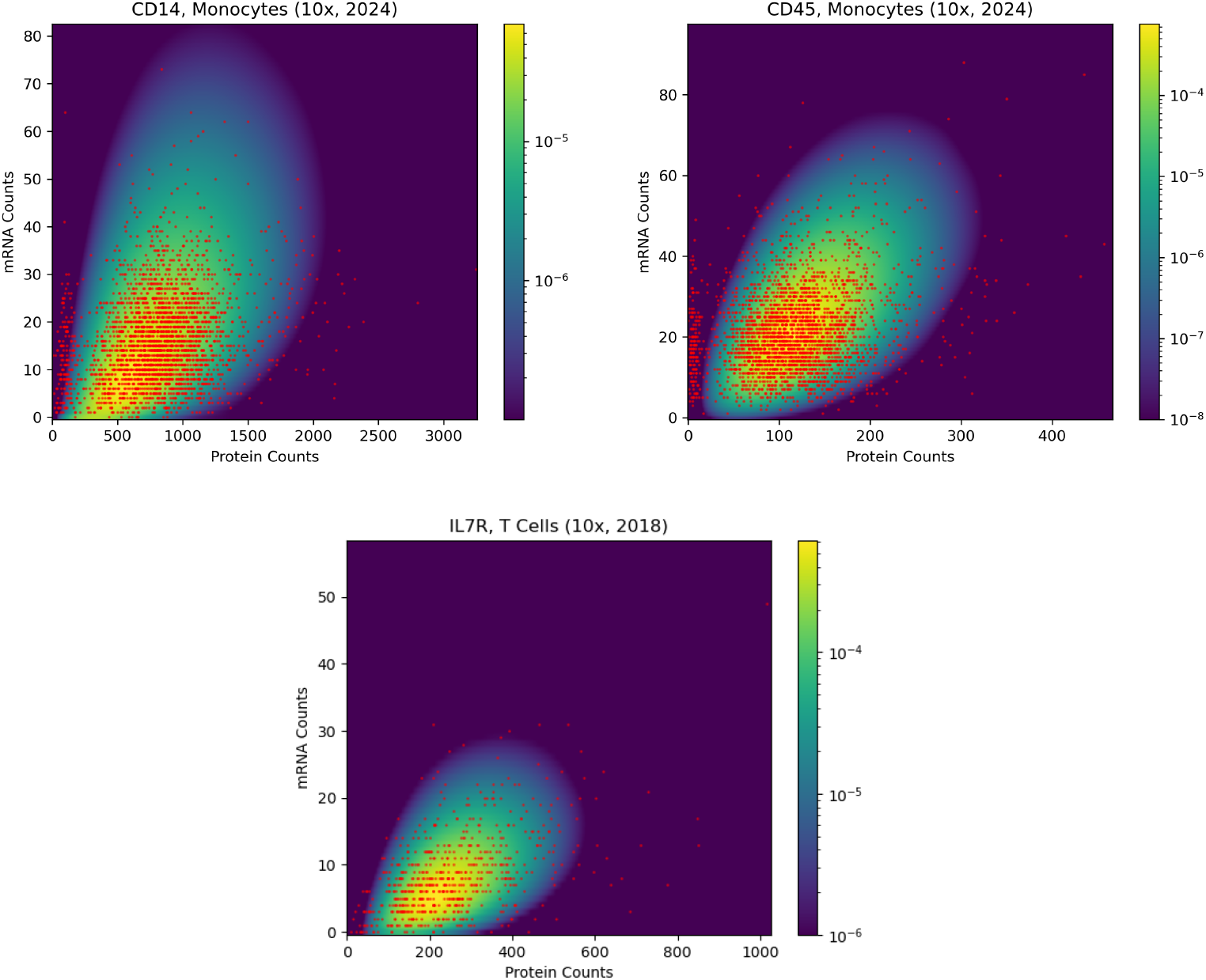
Fits to genes. The color map shows the calculated probability density at the optimal biological parameters, on a logarithmic scale. The red scatter points are the observed distribution of mRNA vs protein counts over cells. The cells were subsetted to monocytes and T cells for the 10x 2024 (10x Genomics 2024) and 2018 (10x Genomics 2018) datasets respectively.

## 3 Discussion

In this work we have presented and explored the power of fitting a joint biophysical model for RNA and protein to single-cell data, and extracted biophysically meaningful parameters. We have demonstrated the robustness of our fitting procedure for the full joint biophysical model via simulation. We have then shown that most of the biophysical parameters can also be inferred to some accuracy using only spliced and protein counts. This approach could be appropriate until the available data includes sufficiently comprehensive unspliced RNA measurements. We have used this reduced model to fit transcriptional rate parameters to CITE-seq data, and have shown that we can successfully reproduce the observed spliced-protein distributions.

Our work highlights the compatibility of current inference frameworks (Gorin et al. 2023) with biophysical models including additional modalities, and we hope that others will continue our approach, perhaps by including chromatin accessibility measurements as an additional model layer. The method described here could also be straightforwardly extended to include principled biophysical clustering using meK-Means (Chari et al. 2023). One limitation of our analysis is the simplification that surface proteins are a proxy for proteins produced within the cell. A non-instantaneous process of surface protein export would add another layer to the model, and the inclusion of protein export is a potential extension of this investigation. Additionally, length-dependent translation (Rogers et al. 2017) could be added to build on the picture provided here. Overall, the approach outlined here represents a first foray into using biophysical modeling to disentangle transcriptional and translational parameters. With development, this method could provide a convenient avenue for determining metabolic rates in systems where direct experimental measurements are difficult to obtain. The full model presented here also provides a framework for the principled integration of single-cell unspliced, spliced, and protein count information. The power of this methodology will increase with the improved reliability of these measurement techniques.

## Data and Code Availability

A github repository with all of the datasets, scripts for fitting parameters, and the simulations presented here is available at: https://github.com/pachterlab/FFP2025.

## Acknowledgments

We thank Andrew LeDuc for helpful discussions. We also thank Charles Trimble for generously funding part of C.F.’s research.

