## Supplementary Information for "Joint Biophysical Modeling of Paired Single-Cell RNA and Protein Measurements"

#### S1 Stationary distribution of mRNA and protein via generating function methods

The generating functions, as defined in Equations 3-4 evolve as follows:

$$\begin{aligned}
\frac{\partial G(z_u, z_s, z_p, t)}{\partial t} &= k(F(z_u) - 1)G + \beta(z_s - z_u)\frac{\partial G}{\partial z_u} + \gamma(1 - z_s)\frac{\partial G}{\partial z_s} \\
&\quad + k_p(z_p - 1)z_s\frac{\partial G}{\partial z_s} + \gamma_p(1 - z_p)\frac{\partial G}{\partial z_p}, \\
\implies \frac{\partial \phi(u_u, u_s, u_p, t)}{\partial t} &= kM(u_u) + \beta(u_s - u_u)\frac{\partial \phi}{\partial u_u} - \gamma u_s\frac{\partial \phi}{\partial u_s} \\
&\quad + k_p u_p(u_s + 1)\frac{\partial \phi}{\partial u_s} - \gamma_p u_p\frac{\partial \phi}{\partial u_p}.
\end{aligned}$$

When $b \to 0, F(z_u) \approx 1 + bz_u$, and this reduces to the constitutive model. For the steady-state solution, we set $\frac{\partial \phi}{\partial t} = 0$ and solve the resulting first order PDE using the method of characteristics (Courant and Hilbert 1962). We write all the variables as functions of $s$, i.e., $\phi(\tilde{u_u}(s), \tilde{u_s}(s), \tilde{u_p}(s), t(s))$, giving the characteristic ODEs:

$$\begin{aligned}
\frac{d\tilde{t}}{ds} &= -1 & \tilde{t}(0) &= t \\
\frac{d\tilde{u}_u}{ds} &= \beta(\tilde{u}_s - \tilde{u}_u) & \tilde{u}_u(0) &= u_u \\
\frac{d\tilde{u}_s}{ds} &= -\gamma\tilde{u}_s + k_p\tilde{u}_p(\tilde{u}_s + 1) & \tilde{u}_s(0) &= u_s \\
\frac{d\tilde{u}_p}{ds} &= -\gamma_p\tilde{u}_p & \tilde{u}_p(0) &= u_p
\end{aligned} \tag{1}$$

and

$$\phi(u_u, u_s, u_p, t) = \phi(\tilde{u}_u(0), \tilde{u}_s(0), \tilde{u}_p(0), 0) + \int_0^t kM(\tilde{u}_u(s))ds. \tag{2}$$

For the bursty model with geometrically distributed burst sizes, we have $M(u) = \frac{bu}{1-bu}$. We have immediately that

$$\tilde{u}_p = u_p e^{-\gamma_p s}, \tag{3}$$

In summary, we need to solve the following:

$$\begin{aligned}
\frac{d\tilde{u}_s}{ds} &= -\gamma\tilde{u}_s + k_p u_p e^{-\gamma_p s}(\tilde{u}_s + 1) & \tilde{u}_s(0) &= u_s \\
\frac{d\tilde{u}_u}{ds} &= \beta(\tilde{u}_s - \tilde{u}_u) & \tilde{u}_u(0) &= u_u \\
\phi(u_u, u_s, u_p, \infty)) &= \int_0^\infty \frac{kb\tilde{u}_u(s)}{1 - b\tilde{u}_u(s)}ds
\end{aligned}$$

### S2 Solving probability distribution numerically

We used Runge-Kutta 4 (Runge 1895) to numerically integrate these equations.

### S3 Biological data analysis

#### S3.1 Data processing

We obtained our sequencing data from the following human PBMC datasets:

- 10k Human PBMCs Stained with TotalSeq™-B Human TBNK Cocktail, Chromium GEM-X Single Cell 3' Universal 3' Gene Expression dataset analyzed using Cell Ranger 8.0.0 (2024, March 13) (10x Genomics 2024)
- 10K Human PBMCs, Gene Expression with a Panel of TotalSeq™-B Antibodies, analyzed using Cell Ranger 3.0.0, (2018, November 19) (10x Genomics 2018)

The raw RNA-seq fastq files were processed with `kb-python` (Sullivan et al. 2025); we used the `nac` workflow with the appropriate 10x technology string. The call for processing the 2024 dataset is shown as an example here:

```
 kb count \
 --overwrite \
 --h5ad \
 --workflow=nac \
 -i /home/cfelce/proMonod/data/ref/human/GRCh38.110/index.idx \
 -g /home/cfelce/proMonod/data/ref/human/GRCh38.110/t2g.txt \
 -x 10xv4 \
 -o /home/cfelce/proMonod/data/RNA_S2 \
-c1 /home/cfelce/proMonod/data/ref/human/GRCh38.110/cdna.txt \
-c2 /home/cfelce/proMonod/data/ref/human/GRCh38.110/nascent.txt \
-m 16G \
--verbose \
--filter bustools \
10k_Human_PBMC_TotalSeqB_3p_gemx_fastqs/gex/
    10k_Human_PBMC_TotalSeqB_3p_gemx_gex1_S2_L001_R1_001.fastq.gz \
10k_Human_PBMC_TotalSeqB_3p_gemx_fastqs/gex/
    10k_Human_PBMC_TotalSeqB_3p_gemx_gex1_S2_L001_R2_001.fastq.gz \
10k_Human_PBMC_TotalSeqB_3p_gemx_fastqs/gex/
    10k_Human_PBMC_TotalSeqB_3p_gemx_gex1_S2_L002_R1_001.fastq.gz \
10k_Human_PBMC_TotalSeqB_3p_gemx_fastqs/gex/
    10k_Human_PBMC_TotalSeqB_3p_gemx_gex1_S2_L002_R2_001.fastq.gz \
10k_Human_PBMC_TotalSeqB_3p_gemx_fastqs/gex/
    10k_Human_PBMC_TotalSeqB_3p_gemx_gex1_S2_L003_R1_001.fastq.gz
10k_Human_PBMC_TotalSeqB_3p_gemx_fastqs/gex/
    10k_Human_PBMC_TotalSeqB_3p_gemx_gex1_S2_L003_R2_001.fastq.gz
10k_Human_PBMC_TotalSeqB_3p_gemx_fastqs/gex/
    10k_Human_PBMC_TotalSeqB_3p_gemx_gex1_S2_L004_R1_001.fastq.gz
10k_Human_PBMC_TotalSeqB_3p_gemx_fastqs/gex/
    10k_Human_PBMC_TotalSeqB_3p_gemx_gex1_S2_L004_R2_001.fastq.gz
```

Some proteins, or protein complexes, corresponded to multiple RNA transcripts in the data. The identifications in Table 1 (10x, 2024) and Table 2 (10x, 2018), were used to map protein counts to RNA counts. The cells were clustered using Leiden clustering (see the scripts at https://github.com/pachterlab/FFP_2025), and subsetted to monocytes for the 10x, 2024 dataset, and to a group of T-cells for the 10x, 2018 dataset.

#### S3.2 Fitted parameters

We fit the bursty transcription with translation model, with Poissonian count sampling, described in the main text, Section 2.1. Note that, since we fit only spliced RNA

| Protein | Subunits (alternate forms) | Ensembls |
|---|---|---|
| CD3 (complex) | CD3$\gamma$ (x1)<br>CD3$\delta$ (x1)<br>CD3$\epsilon$ (x2) | ENSG00000160654<br>ENSG00000167286<br>**ENSG0000019885** |
| CD4 | - | ENSG00000010610 |
| CD8 | CD8a<br>(CD8b) | ENSG00000153563<br>(ENSG00000172116) |
| CD11C | - | ENSG00000140678 |
| CD14 | - | ENSG00000170458 |
| CD16 | CD16a<br>(CD16b) | ENSG00000203747<br>(ENSG00000162747) |
| CD19 | - | ENSG00000177455 |
| CD56 | - | ENSG00000149294 |
| CD45 | - | ENSG00000081237<br>(ENSG00000262418) |

**Table 1** Proteins (complexes) with their constituent subunits or alternative forms, and the corresponding Ensembl IDs, for the 10x 2024 dataset. Where alternate forms are given in parentheses, the non-parenthesized Ensembl ID was used. Where subchains of a complex are shown, the bolded ensembl was used.

| Gene Symbol | Ensembl ID |
|---|---|
| PTPRC | ENSG00000081237 |
| FCGR3A | ENSG00000203747 |
| CD247 | ENSG00000198821 |
| CD3E | ENSG00000198851 |
| CD3D | ENSG00000167286 |
| CD3G | ENSG00000160654 |
| FUT4 | ENSG00000196371 |
| NCAM1 | ENSG00000149294 |
| CD4 | ENSG00000010610 |
| IGHG1 | ENSG00000211896 |
| IGHG2 | ENSG00000211893 |
| ISG20 | ENSG00000172183 |
| CD19 | ENSG00000177455 |
| PDCD1 | ENSG00000188389 |
| CD8A | ENSG00000153563 |
| TIGIT | ENSG00000181847 |
| IL7R | ENSG00000168685 |
| CD14 | ENSG00000170458 |

**Table 2** Final one-to-one mapping, used for the 10x 2018 dataset, between protein (gene) symbols and Ensembl IDs.

and protein counts, the technical sampling rate for unspliced counts only affected the method of moments initialization values.

We show the optimal biological parameters for each fit in Table 3, along with the technical parameters used (capture rates $\lambda_{u,s,p}$ for unspliced, spliced and protein counts respectively), and the initialization method used for fitting.

| | **2024, CD14** | **2024, CD45** | **2018, IL7R** |
|---|---|---|---|
| **Cell Type** | Monocytes | Monocytes | T-cell cluster |
| $\log_{10} b$ | 3.04 | 1.96 | 4.2 |
| $\log_{10} \beta$ | -0.403 | -0.993 | -0.665 |
| $\log_{10} \gamma$ | 1.77 | -0.406 | 1.31 |
| $\log_{10} k_p$ | 3.5 | -1.14 | 1.24 |
| $\log_{10} \gamma_p$ | -0.634 | -0.862 | -0.764 |
| $\log_{10} \lambda_u$ | 0.0 | - | 0.0 |
| $\log_{10} \lambda_s$ | 0.0 | -1 | -2.0 |
| $\log_{10} \lambda_p$ | -2.5 | 0 | -2.5 |
| **Initialization** | Moments estimate | 10 random restarts | Moments estimate |

**Table 3** Optimal biophysical parameters for each of the fits shown in the main text, and the corresponding technical capture rates. The cell type and initialization method for the fit are also given.
